# Variability of soybean response to rhizobia inoculant, Vermicompost, and a legume-specific fertilizer blend in Siaya County of Kenya

**DOI:** 10.1101/476291

**Authors:** Catherine Mathenge, Moses Thuita, Cargele Masso, Joseph Gweyi-Onyango, Bernard Vanlauwe

## Abstract

Rhizobia inoculation can increase soybean yield, but its performance is influenced by soybean genotype, rhizobia strains, environment, and crop management among others. The objective of the study was to assess soybean response to rhizobia inoculation when grown in soils amended with urea or Vermicompost to improve nitrogen levels. Two greenhouse experiments and one field trial at two sites were carried out. The first greenhouse experiment included soils from sixty locations, sampled from smallholder farms in Western Kenya. The second greenhouse experiment consisted of one soil selected from soils used in the first experiment where inoculation response was poor. The soil was amended with Vermicompost or urea. In the two greenhouse experiments, Legumefix® (inoculant) + Sympal (legume fertilizer blend) were used as a standard package. Results from the second greenhouse experiment were then validated in the field. In the first greenhouse trial, soybean response to inoculation was significantly affected by soil fertility based on nodule fresh weight and shoot biomass. Soils with low nitrogen had low to no response to inoculation. After amendment, nodule fresh weight, nodule effectiveness, nodule occupancy, and shoot dry biomass were greater in the treatment amended with Vermicompost than those amended with urea (Legumefix® + Sympal + Vermicompost and Legumefix® + Sympal + urea). Under field conditions, trends were similar to the second experiment for nodulation, nodule occupancy, and nitrogen uptake resulting in significantly greater grain yields (475, 709, 856, 880, 966 kg ha^−1^) after application of Vermicompost at 0, 37, 74, 111, and 148 kg N ha^−1^, respectively. It was concluded that soybean nodulation and biological nitrogen fixation in low fertility soils would not be suppressed by organic amendments like Vermicompost up to 148 kg N ha^−1^.

## Introduction

Soybean (*Glycine max* L. Merr) is one of the world’s most important legumes in terms of production and trade and has been a dominant oilseed since the 1960s [1]. The crop is well known for its high protein content (about 40%) [2]. Additionally, it can improve soil properties and soil biological health by soil nitrogen enrichment through N_2_ fixation and subsequent mineralization of shoot and root biomass [3]. It therefore represents a significant opportunity in sub-Saharan Africa (SSA), where over 80% of the soils are nitrogen deficient [85], and over 39% of the children under 5 years are stunted because of malnutrition caused by nutrient deficiency, particularly proteins [4], contributing to over one third of child deaths [5]. Integration of soybean in smallholder farming systems would thus not only improve human nutrition when the crop is included in diets but also soil productivity. Such benefits would materialize when good agronomic practices, including integrated soil fertility management, are implemented in soybean production systems.

Crop production, including soybean, faces several constraints which include abiotic and socioeconomic factors accounting for production discrepancies across regions in SSA. Consequently, grain yields remain low compared to other regions in the world [6]. Integrated soil fertility management (ISFM), has been proposed as a viable way towards the sustainable intensification of smallholder agriculture [7]. The high cost of inputs for nutrient replenishment or soil amendment has however limited their adoption by resource-constrained smallholder farmers [8]. Utilization of soybean varieties with high biological nitrogen fixation (BNF) potential and application of rhizobia inoculants would represent a cost-effective option to reduce mineral N application [9-14). Studies on N_2_ fixation in soybean using different methodologies revealed that soybean shows a strong demand for nitrogen, up to 80 kg N per 1000 kg of soybean grain for optimal development and grain productivity [15, 16]. Soybean can fix N from the atmosphere ranging from 0 to 450 kg N ha^−1^ [17, 18]. Under environments conducive for N fixation, over 60 to 70% of the N requirement of the soybean can be derived from BNF [19], while the balance could be derived from the soil N stock. Conversely, it has been reported that BNF could be as low as 5 kg N ha^−1^ in depleted soils, which are quite common in the smallholder farming systems in SSA, which would imply reliance on nitrogen fertilizers even for legume crops [20].

Low soil fertility in SSA is often characterized by low available phosphorous (P), nitrogen (N), organic matter (C_org_), and soil acidity, among others [21]. Such parameters must be corrected as they are an integral part of the interaction of legume genotype, rhizobia strain, environment, and crop management, which determines the performance of BNF in particular and legume productivity in general [22-25]. Soil organic carbon is a key driver of soil fertility that could even impede the performance of non-limiting factors, when it is below a certain level in a specific soil type [26]. Response to inorganic fertilizers could be enhanced by the addition of organic matter [27]. However most agricultural soils in SSA contain low levels of organic carbon due to competing use of organic residues [28, 29]. Initiatives that promote rhizobia inoculation in legume production in Africa generally recommend the application of nutrients such as P, and lately, more balanced blends have been developed for use with inoculums but do not include N [14, 23, 30, 31]. This is due to the general assumption that rhizobia would supply the N required by the legume and applying mineral N would inhibit nodulation. While such inhibition has been well-documented [32], this could be different in low fertility soils that are N deficient [33].

Starter N is sometimes needed to achieve a substantial yield of legumes including soybean when the symbiotic N_2_ fixation is unable to provide enough nitrogen [34].

The objective of the study was thus to assess whether soils with a low inoculation response could be improved by amendment. It was hypothesized that an organic amendment would perform better than a mineral N fertilizer, given the expected high correlation between organic carbon and total nitrogen in agricultural soils [35].

## Materials and methods

### Characterization of the study soils

Two greenhouse experiments were established at the International Centre of Insect Physiology and Ecology (*icipe*), Duduville campus, Nairobi, Kenya. Soils were collected from sixty farms of Siaya County where low soybean response to inoculation was observed [25, 84] (Fig 1) (where a varied response to an ISFM soybean package had been observed) at a depth of 0‒20 cm, air dried, and thoroughly mixed to pass through a 2-mm sieve. Subsamples were analyzed for physical, chemical, and microbiological properties prior to planting. The soils parameters analyzed were organic Carbon determined by chromic acid digestion and spectrophotometric analysis [37], total N (%) determined from a wet acid digest [38], and N analyzed by colorimetric analysis [39]. Soil texture was determined using the hydrometer method; soil pH in water determined in a 1:2.5 (w/v) soil: water suspension; available P using the Mehlich-3 procedure [40] and the resulting extracts analyzed using the molybdate blue procedure [41]; and exchangeable cations (Ca, Mg, and K) extracted using the Mehlich-3 procedure and determined by atomic absorption spectrophotometry. Estimation of rhizobia in the soils was done using the most probable number count [42]; soybean variety TGx1740-2F was used as a trap crop grown in N free and autoclaved sterile sand.

### Greenhouse experiments

The first greenhouse experiment was laid as a Completely Randomized Design (CRD) including: (i) 60 soils collected from the sites indicated in Fig 1 with N and C_org_ ranges of 0.029‒0.21% and 0.53‒2.1%, respectively, (ii) two treatments, i.e., with and without inoculation (Legumefix® + Sympal) replicated 3 times for a total number of 360 experimental units. Co-application of Legumefix® and Sympal, as an inoculation package, was informed by previous findings [23, 36]. Sympal is a legume-specific fertilizer blend (N: P_2_O_5_: K_2_O 0: 23:15 + 10CaO + 4S + 1MgO + 0.1Zn) and was applied at a rate equivalent to 30 kg P ha^−1^ and thoroughly mixed with the soil for the inoculated treatments (i.e.,≈300 kg Sympal ha^−1^). Soybean variety (TGx1740-2F) was selected due to its better nodulation with a range of rhizobia than local varieties in different parts of Kenya [43]. Seeds were surface-sterilized by soaking in 3.5% NaClO solution for 2 min and rinsed thoroughly 5 times with sterile distilled water. Soils were weighed to fill perforated 2.5-kg pots. Legumefix® for soybean (containing *Bradyrhizobium japonicum* strain 532c) from Legume technology Inc (UK) was used at a rate of 10 g per kg soybean seeds for the inoculated treatments. Three healthy seeds of uniform size were then planted per pot and thinned to one plant per pot of comparable height and vigor at 2 weeks after planting. Routine management practices such as watering were carried out till termination of the experiment, i.e., at 50% podding. This trial was thus intended to determine soybean response to co-application of inoculation and Sympal in various soils characterized by a gradient of nitrogen content.

In the second greenhouse experiment, one of the 60 experimental soils (Trial Site 17 in Fig1), that showed low response to inoculation in the 1^st^ greenhouse experiment based on low nodule fresh weight and shoot dry weight observed was amended either with Vermicompost (Phymyx) or urea. Vermicompost was chosen as a slow release form of N compared to urea. A slow N release would reduce the negative effect of N application to nodulation at the early growth stages of soybean. The soil was collected from an area of 4 × 3 m at a depth of 0‒20 cm and homogenized after air drying and sieving. Vermicompost (Vc) was applied at 5 levels with even intervals including a control (at rates equivalent to 0, 2.5, 5, 7.5, and 10 t Vc ha^−1^). Equivalent amounts of N were applied using urea (46% N). The rates of N were thus 0, 37, 74, 111, and 148 kg N ha^−1^. Selected chemical properties of the batch of the Vermicompost used in this study based on the product analysis were: total N (0.88%), organic C (7.31%), available P (0.39%), Ca (0.29%), Mg (0.1%), K (0.22%), in addition to a pH that was approximately neutral (6.7%). It was also expected to contain trace micronutrients (not determined) and is made by composting plant residue and livestock manure. The Legumefix® for soybean inoculant was used at the same rate as the 1^st^ greenhouse experiment. The trial was laid as a CRD and each treatment replicated 3 times for a total of 60 experimental pots. Planting, management, and harvesting were done as described in the 1^st^ greenhouse experiment. The trial was thus intended to determine whether application of starter N would improve soybean response to co-application of inoculant and Sympal in a soil with both low nitrogen levels and response to inoculation, and whether there was a systematic difference between Vermicompost and urea as sources of N.

### Field trial

The field trial was intended to validate the findings of the second greenhouse trials in field conditions with a focus on the best performing source of N and determine the yield performance. It was conducted at trial site 17 (Soil B) and site 7 (Soil A) (Fig 1). Site 17 was the farm at which soil was collected for the second greenhouse experiment. Soils from both sites had similar response trends in nodulation and shoot biomass as the first greenhouse experiment even though they did not have the same physical chemical characteristics (Table 1) and thus were chosen for the field trial validation. The field trial was conducted during the long rains (April to August) of the 2016 cropping season. The treatments at each site were laid out in a full factorial randomized complete block design (RCBD) where Sympal was applied at the rates used in the greenhouse trials (0 and 30 kg P ha^−1^). The five rates of Vermicompost used in the second greenhouse experiment were applied, i.e., equivalent to 0, 2.5, 5, 7.5, and 10 t ha^−1^, whereas inoculation was done using Legumefix® for Soybean at the same rate as the greenhouse trials. The maximum of 10 t ha^−1^ was based on the general recommendation for compost application in the region. The plot sizes were 3 m × 3 m with a 0.5 m alley between the plots and 1 m between the three blocks. Soybean was planted at a spacing of 50 cm (between rows) × 5 cm (within rows) at the onset of the long rainy season (April 2016). Vermicompost and Sympal were applied in furrows and mixed with soil before placement of seeds to avoid direct contact with the seed. Seed sterilization and inoculant application rates were as used in the greenhouse trials. The trials were kept weed free by mechanical weeding.

**Table 1.**
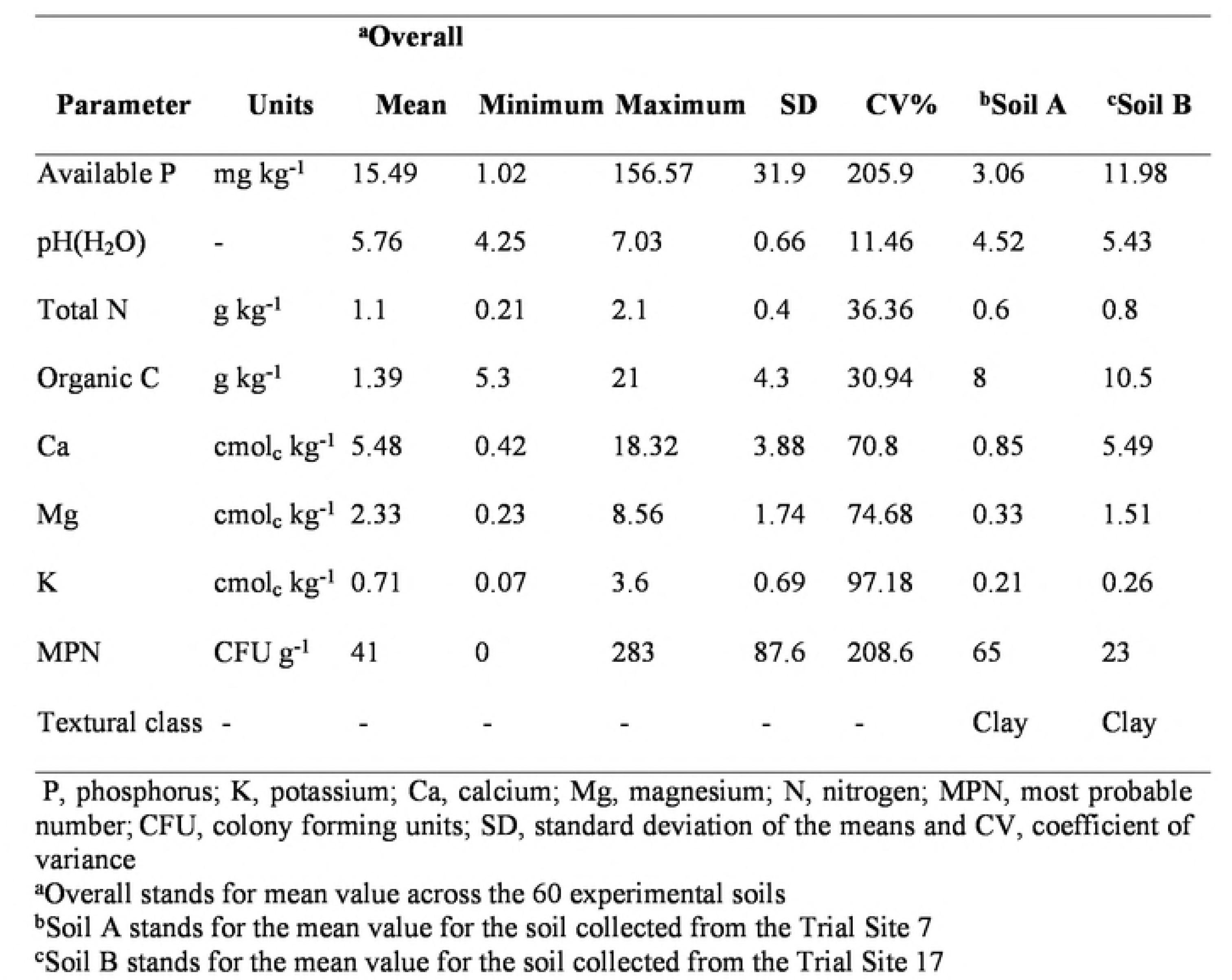
Selected chemical, microbiological, and physical properties of the sixty experimental soils with details of Trial site 7 (Soil A) and Trial site 17 (Soil B)

### Data collection

In the greenhouse experiments the plants were harvested at 50% podding. Shoots were cut using a clean, sharp knife at 1 cm above the soil surface. The pots were emptied into a 2-mm sieve and soil washed to isolate the nodules from the roots. Nodule fresh weight, shoot biomass, and nodule occupancy were captured in both greenhouse experiments, whereas in the second greenhouse experiment additional data collected were nodule effectiveness and N uptake.

Fresh nodules were surface sterilized and stored in glycerol for nodule occupancy determination. Nodule occupancy was then done using the Polymerase chain reaction-Restriction fragment length polymorphism (PCR-RFLP) method. This involved amplification and restriction of the 16S-23S rDNA intergenic spacer region. A maximum number of eight nodules from each of the three replicates per treatment (24 nodules) were crushed separately in 150 µl of sterile water and DNA extracted [44]. Amplification of DNA (PCR) was conducted using rhizobia specific primers [45, 46]. Due to the low number of nodules in the low rates of Vermicompost and urea treatments, only the three upper rates and their respective combinations (74, 111, and 148 kg N ha^−1^) were considered. In addition, restriction was only conducted for PCR products of a single band of 930‒1050 bp with restriction endonucleases *Moralla species* (*Msp* I). The stain with identical fragment size and number were classified into the same profile and the profiles used to score the inoculant (Legumefix® for soybean) efficacy in percentages [13].

Nodule effectiveness was carried out [47]. Fresh shoots were dried at 60 °C until constant weight (approximately 48 hours) to obtain the dry weight. The shoots were later milled for total N analysis by the modified Kjeldahl method. Nitrogen uptake at 50% podding was determined as the product of shoot dry biomass and the respective nitrogen content in the shoot and reported as g N plant^−1^.

In the field trial, the parameters recorded at 50% podding were nodule fresh weight, nodule effectiveness, nodule occupancy, shoot dry biomass, and N uptake, while at harvest grain yield was determined. Eight to ten plants were taken from one of the inner rows about 50 cm from the beginning of the line at 50% podding. Nodules were dug out and washed for nodule fresh weight determination and shoots collected for drying and weighing. A sample, i.e., 10% of the total number of nodules counted per treatment, was taken and used for determining nodule effectiveness. At physiological maturity, when 95% of the pods had turned golden yellow, all plants were harvested from the net plot excluding the outer rows. Number and weight of all plants were recorded from each plot and grains and haulms separated and weighed. The grains were later oven-dried to a constant weight.

### Data analysis

In the two greenhouse trials and the field trial, the analysis of variance (ANOVA) was conducted to assess the effects of the various sources of variation, i.e., treatments using SAS version 9.4. The effects of the various factors and their interactions were assessed using standard error of difference (SED) on the mean. The significance level of the models was set at p < 0.05. In the first greenhouse experiment, box-and-whisker plots were also used to summarize the information on nodule fresh weight and shoot biomass given the large number of experimental soils (sixty data points). The assessment of nodule occupancy for the greenhouse and field trials was based on profiles with similar bp fragments in size after restriction and compared to the IGS profile of strain *B. japonicum* 532c and converted to a percentage for each IGS profile group.

## Results

### Soil properties

Selected soil properties of the experimental soils including details of trial sites 7 and 17 before the beginning of the greenhouse and field trials are presented in Table 1. A wide variation in soil properties was noted with coefficients of variance (CV) ranging from 11 to 208% with most of the parameters falling under what is considered low. There was a strong correlation between total N and C_org_ (r = 0.94). Rhizobia population for the selected soils ranged from 0 to 2.83 × 10^2^ CFU g^−1^ of soil; hence, the response to inoculation was expected.

### Nodulation

In the first greenhouse experiment, the nodule fresh weight (NFW) significantly varied across soils irrespective of inoculation, which was related to the wide variation in soil properties (Table 2; Fig 2). On average inoculated (with) plants had higher NFW than uninoculated (without) plants (Fig 2). The NFW was generally low in the soils of low fertility, which calls for further investigation to reduce the spatial variability.

In the 2^nd^ greenhouse experiment, co-application of starter N in the form of Vermicompost or urea (at low rates), inoculation, and Sympal significantly increased nodule fresh weight (p < 0.05) (Table 2). The upper rate of urea led to a decrease in NFW contrary to Vermicompost, which could be related to the difference in the availability of N from the two sources. Vermicompost co-applied with inoculation and Sympal consistently recorded a significantly higher nodule fresh weight than urea co-applied with inoculation and Sympal (Fig 3a).

In field conditions, NFW was improved by inoculation at Trial site 17 compared to Trial site 7 (Fig 3b), which could be related to the initial fertility level of the sites (Table 1). Conversely, in the absence of inoculation, Trial site 7 performed better than Trial site 17 (Fig 3b), which could be associated with the abundance of soybean nodulating rhizobia at Trial site 7 (Table 1). Regardless of the sites and inoculation, application of Sympal (Fig 3c) and Vermicompost (Fig 3d) improved NFW, which implied that the nodulation of soybean by native rhizobia could be improved with good soil fertility management.

### Nodule effectiveness

In the 2^nd^ greenhouse experiment, N-amendment using Vermicompost or urea co-applied with inoculation and Sympal significantly increased the percentage of effective nodules (p < 0.05) (Table 2; Fig 4a). Vermicompost co-applied with incoulation and Sympal consistently had a higher percentage of effective nodules compared to urea, inoculation and Sympal (Fig 4a). In field conditions, inoculation at Trial Site 7 and Trial Site 17 improved nodule effectiveness at both sites, but co-application with Sympal showed better performance at Site 17 than Site 7 when compared to inoculation without Sympal (Fig 4b). Conversely, in the absence of inoculation, Sympal improved nodule effectiveness at Site 7 more than Site 17, but when Sympal was not applied, nodule effectiveness was similar at both sites (Fig 4b). While Sympal contributed to the improvement of nodule effectiveness, the magnitude of the response demonstrated that inoculation was very critical to enhance the percentage of effective nodules. This suggested the introduced strains not only increased the abundance of rhizobia cells in the rhizosphere, but were also effective in field conditions. Significant improvement of nodule effectiveness following Vermicompost application was made at a total N rate ≥ 74 kg ha^−1^ irrespective of inoculation and Sympal at both sites (Fig 4c).

### Nodule occupancy

For nodule occupancy, IGS profiles as a function of total number of nodules with PCR-RFLP (930‒1050 bp) bands was used. In the 1^st^ greenhouse experiment, three IGS profile groups were obtained from PCR-RFLP analysis. The IGS profile I (91%) (inoculant strain) was dominant in the inoculated soils, while IGS profiles I and III, were in almost equal proportion in unioculated soils (46 and 40%) respectively (Table 3). In the 2^nd^ greenhouse experiment, nodule occupancy by the inoculant strain consistently increased with the increase in Vermicompost rates in the uninoculated treatment, showing that the strain in the rhizobia inoculant is present in the study region due to a previous history of soybean cultivation with the inoculant strain in the two sites (Table 4). An increased rate of N from Vermicompost up to 148 kg ha^−1^ did not suppress nodule occupancy by the inoculant strain, while at a rate of 148 kg N ha^−1^ urea nodulation was suppressed to the extent that no nodules were found, with and without inoculation. This could be related to the slow release of N in Vermicompost compared to urea. For the rates of 74 and 111 kg N ha^−1^, under co-application of the rhizobia inoculant and Sympal, all the nodules analyzed carried the inoculant strain. Based on the results reported in Fig 3a (nodule fresh weight) and Fig 4a (nodule effectiveness) at 148 kg N ha^−1^ from urea, it is likely that some native strains that can nodulate soybean were not detected by the specific primers used to assess the nodule occupancy and thus total number of nodules analyzed were not equal in all the treatments. This often occurs when some bacteria have acquired genes that enable nodulation but may not have all the required genes to allow detection by the set of primers; further investigation would be required.

In the field trial, the highest inoculant strain recovery was observed with the combination of inoculation, Vermicompost, and Sympal demonstrating the relevance of the combination to supply additional nutrient, organic matter, and rhizobia particularly in the low fertility soil at Site 7 (Table 4). The highest inoculant strain recovery was attained when Vermicompost was applied at 74 kg N ha^−1^ and 111 kg N ha^−1^ and combined with Sympal and inoculation at Site 7 (94%), while there was a slight reduction at 148 kg N ha^−1^ for the same inputs though the inoculant strain recovery was still higher than 66% (Table 4). At Site 17, which was slightly more fertile than Site 7, the value addition of co-applying inoculation and Sympal in the presence of Vermicompost was reduced, except at 111 kg N ha^−1^ (Table 4). In the absence of the rhizobia inoculant, co-application of Vermicompost and Sympal did enhance the nodule occupancy by native strains other than the inoculant strain, which could be less effective based on the results on nodule effectiveness (Fig 4b). In general, a consistently higher percentage of nodules occupied by the inoculant strain was observed in the inoculated and amended soils for both greenhouse and field conditions at moderate levels of N (74 and 111 kg N ha^−1^ regardless of the source of N). This suggests that the introduced strain was more competitive in the amended soils and explains the higher percentage of effective nodules (Fig 4b). The recovery of the inoculant strain from the uninoculated treatments (especially in the 2^nd^ greenhouse experiment) was attributed to the previous history of soybean cultivation with the same inoculant in the two farms.

### Shoot biomass

On average, the inoculated treatment gave a higher shoot dry weight than the uninoculated soils in the 1^st^ greenhouse trial, with an increase of 38% over the control (Fig 2), but the improvement of shoot biomass following inoculation significantly varied across soils (Table 2). In the 2^nd^ greenhouse experiment, co-application of N amendments (Vermicompost or urea) with inoculation and Sympal enhanced shoot dry biomass (Fig 5a). When N was applied as Vermicompost, the value addition of inoculation and Sympal was found at the low rate of N (equivalent to 37 kg N ha^−1^) and in the untreated control (no N). Conversely, when N was applied as urea, the value addition of inoculation and Sympal was found across the five rates of N. The difference between the two sources of N can be related to the additional nutrients in Vermicompost compared to urea that only supplied N. Across treatments, the highest shoot dry biomass at 50% podding was found at 148 N kg^−1^ applied as urea and combined with inoculation and Sympal. This could be attributed to the fact that nitrogen from urea was readily available for uptake and resulted in vigorous vegetative growth and more biomass accumulation at the early stage of the crop with minimal N losses in greenhouse conditions.

In field conditions, co-application of Vermicompost and inoculation significantly improved shoot dry biomass compared to Vermicompost in the absence of inoculation, particularly when Sympal was not applied (Fig 5b). When Sympal was added to both combinations (Vermicompost with and without inoculation), the difference in shoot dry biomass was reduced, which could be related to improved utilization of N when other limiting nutrients are added. On average, the shoot dry biomass was higher at Site 17 than Site 7 irrespective of the treatments (Fig 5c), which was consistent with the initial chemical properties of the two sites (Table 1).

### Shoot biomass N uptake

In the 2^nd^ greenhouse experiment, co-application of Vermicompost or urea as a source of starter N with inoculation and Sympal significantly increased biomass N uptake when compared to the starter N sources in the absence of inoculation and Sympal (Table 2; Fig 6a). When Vermicompost or urea was not co-applied with inoculation and Sympal, increased rates of Vermicompost enhanced biomass N uptake, while increased rates of urea reduced N uptake. This could be related to the improved soybean growth in the presence of Vermicompost related to the additional nutrients in the inputs, which would have improved the root system development (data not collected) and consequently N uptake. Application of inoculation and Sympal to both starter N treatments further enhanced plant development and therefore N uptake.

In the field conditions, co-application of Vermicompost and Sympal enhanced N uptake at Site 7 (which was less fertile) more than Site 17 (Fig 6b). In the absence of Sympal, N uptake was similar at both sites when Vermicompost was applied. On average, rhizobia inoculation showed higher N uptake than uninoculated plants, irrespective of trial sites, Vermicompost, and Sympal (Fig 7c).

### Grain yield

When soybean was inoculated, yields were higher at Site 17 than Site 7, while both sites had similar yields in the absence of inoculation (Fig 7a). Hence, the apparent difference in soil fertility at the two sites (Table 1) was not enough to show a difference in yields without soil amendment. Amendment with Vermicompost increased soybean grain yield from a rate of 74 kg N ha^−1^ compared to the absolute control (Fig 7b); this rate of N was equivalent to five tons of Vermicompost ha^−1^. Grain yields significantly increased on amendment (475, 709, 856, 880, 966 kg ha^−1^) after application of Vermicompost at 0, 37, 74, 111, and 148 kg N ha^−1^, respectively. All the measured parameters reported correlated significantly to grain yields particularly at Site 7 (data not shown), which showed that amending low fertility soils using various combinations of inputs like rhizobia inoculant, Sympal, and Vermicompost could enhance soybean growth and yield assuming no other limiting factors.

## Discussion

In this study, overall the effects of four key factors: site (soil), rhizobia inoculant, starter N (Vermicompost or urea), and a legume-specific fertilizer blend (Sympal) and their interactions on soybean productivity traits including nodulation, nodule effectiveness, nodule occupancy, shoot dry weight, N uptake, and yield were evaluated. These productivity traits were improved by various combinations of the three inputs, but in most cases, there was a significant site or soil effect. Previous studies demonstrated that legume response to inoculation is generally affected by (i) legume genotype, (ii) rhizobia strain, (iii) environments like soil fertility, soil amendment, and water management, and (iv) crop management such as weeding, spacing, and pest and disease control [23, 48]. In this study, the focus was on aspects related to soil fertility improvement to enhance soybean productivity traits. The hypothesis that starter N, particularly in its organic form, would improve soybean response to rhizobia inoculants and legume-specific fertilizer blends (without N) in low fertility soils was confirmed and it is crucial to understanding the underlying mechanisms.

### Need for starter N to improve soybean response to inoculation in low fertility soils

The soils used in the three experiments were low in nitrogen levels as reported [36]. Nitrogen is a major limiting factor in plant growth and development. In low fertility soils, there is a need to explore various nutrient replenishment avenues to establish best practice management options for improved soybean response to inoculation [23]. In soils with low nitrogen, a moderate amount of “starter nitrogen” would be required by the legume plants for nodule development and root and shoot growth before the onset of BNF [49, 50]. In the low N soil used in the second greenhouse experiment, amendment with two nitrogen sources (Vermicompost and urea) significantly increased soybean productivity traits suggesting the nitrogen supplied played a great role in soybean growth before a symbiotic relationship of the host crop and rhizobia was fully functional. Although insignificant responses of starter N have been reported [51], positive responses have been reported by several studies which demonstrates the need of starter N, particularly in low fertility soils as it was the case in this study [52-56]. There is need to determine the threshold value of soil N content (% or g N kg^−1^ soil) above which, starter N would not be required.

### Preference of an organic source for starter N in low fertility soils

The Vermicompost treatments performed better in all the measured parameters compared to the urea treatments. Although N supplied by urea was readily available for the plant uptake, N alone could not explain the significant increase in the soybean growth traits observed. Vermicompost not only was a source of slow-release N, but also other essential nutrients such as Ca, Mg, and K, which are essential for optimal plant growth. Organic sources of N also improve soil organic carbon, which has a significant effect on soil fertility including rhizobia survival [57]. In general, soil total N and organic matter are highly correlated as found in this study. In low organic matter soils, organic amendments act as a source of nutrients, improve soil structure, and increase biodiversity and activity of the microbial population [58, 59]. Use of organic amendments to improve nutrient-depleted soils in SSA in general and western Kenyan in particular [23] would improve the physical, chemical, and biological characteristics of soil [58]. This implies that soil amendment with Vermicompost, or similar organic inputs, would be a good practice to improve soybean response to inoculation as nodulation and nodule effectiveness were not suppressed up to a rate of 148 kg Vermicompost-N ha^−1^. Furthermore, use of organic amendments including organic fertilizers in integrated soil fertility management to supply both nutrients and organic matter would be more conducive to sustainability and resilience of the cropping systems than sole application of inorganic fertilizers.

### Balanced fertilization to improve soybean response to inoculation

Significant variation of soybean response to rhizobial inoculation was observed across the sixty soils in greenhouse conditions, which was validated in field conditions at two sites. Success of soybean rhizobia inoculation is dependent on soil fertility and site location [59]. Based on recommendations [60] and the soil analysis results, the study soils from sixty locations in western Kenya had very low to moderate fertility, which agreed with earlier report [23]. This wide variation in soil properties with most of the parameters falling under low to very low [61, 62] could explain the variation of the soybean response to inoculation. Similar findings of spatial variation of soybean response to biological inoculants across locations was previously reported [36]. Edaphic factors such as nutrient P and N availability and soil pH determines the effectiveness of inoculant used [36]. This has also been confirmed in our ongoing investigation on the effect of soil acidity and liming on soybean productivity traits under inoculation (unpublished). Soil amendment to improve the fertility including balanced fertilization is therefore crucial to reduce the spatial variability of soybean response to inoculation, assuming no other limiting factors.

In field conditions, nodule fresh weight and effectiveness were improved by the application of Sympal and/or Vermicompost. Shoot dry weight was enhanced by co-application of Vermicompost, Sympal, and inoculation, while a combination of Vermicompost and Sympal increased biomass N uptake and Vermicompost boosted grain yield. This was in line with previous findings [ 6, 63-66]. Soil amendment improved the effectiveness of the nodules and the competitiveness of the introduced strain to occupy a significant number of nodules, as shown by the nodule occupancy. Vermicompost and Sympal contained various nutrients including macro-, secondary, and micronutrients, which are essential to plant growth and effective nodulation. A package of fertilization interventions based on proper soil fertility diagnosis in legume cropping systems including organic inputs, a legume-specific fertilizer blend conducive to nodule formation, and efficacious rhizobia inoculants would be more effective than a sole application of one component of the package [65, 67-70]; though profitability analysis would be required to inform the choice of package to recommend. Hence, current development initiatives that promote rhizobia inoculation without necessary soil fertility diagnosis or only focus on co-application of phosphorus and rhizobia inoculants must be revisited to consider balanced fertilization. Effective legume rhizobia inoculation only adds N in the cropping systems so there is a need to ensure that the other nutrients are available at appropriate levels for optimum plant growth. Availability of essential nutrients and moderate levels of nitrogen generally enhance nodule formation and functioning [71, 73]. High rates of nitrogen fertilizers however have been shown to inhibit nodule formation in both controlled and field conditions [24, 34, 74, 75]. Hence, investigations to determine the threshold values, depending, among others, on soil types, below which starter N would be required to improve legume response to inoculation in low fertility soils, are needed.

### Effectiveness of inoculant rhizobial strains

Response to rhizobia inoculation is expected in soils of low native rhizobia or where the compatible rhizobia of the host legume are absent [76, 77]. The rhizobia populations in the sixty soils were below 1.0 × 10^3^ CFU g^−1^ of soil, which has been reported as the minimal population of native rhizobia for a response to inoculation to be achieved for legume crops like soybean [78]. The capacity of an inoculant strain to occupy nodules on the host depends on environment factors such as the presence of indigenous rhizobia and soil type [79, 80]. The increased nodule weight and shoot biomass over the control due to rhizobia inoculation indicated that the introduced strain was more effective than the indigenous bradyrhizobia. This was in line with previous studies [13, 56, 81-83] which reported significant increases in nodulation and biomass with rhizobia inoculation. The soybean increased biomass, nodulation, and effective nodules due to inoculation confirms the need to inoculate soybean seeds in the soils of the selected sites. Even though the variety TGx1740-2F is promiscuous, nodule occupancy analysis confirmed successful inoculation. Inoculation with Legumefix® for soybean significantly increased the percentage of effective nodules and nodule occupancy both in greenhouse and field experiments. Nodule effectiveness and occupancy are important indicators of efficient soybean rhizobia symbiosis [47, 80]. The yield increase following inoculation at both sites was in line with other reported findings [14, 65, 81, 87]. As mentioned above, to optimize soybean response to rhizobia inoculants, soil amendment with organic sources of nutrients and legume-specific fertilizer blends in low fertile soils will be of great importance not only in Siaya County of Kenya, but also across SSA where nutrient depletion is widely spread [4, 86] in addition to address issues related to factors like legume genotype, efficacy of rhizobia strains, as well as good crop and water management.

## Conclusion

Soil amendment with Vermicompost, inoculation, and Sympal in low fertility soils increased soybean productivity traits including yields. Soybean response to inoculation was affected by soil properties. Vermicompost supplied both nutrients and organic carbon, while Sympal contributed additional nutrients, which improved the nutrient status of the low fertility soils and consequently soybean response to inoculation. Development initiatives focusing on legume inoculation or co-application of rhizobia inoculants and phosphorus fertilizers only, without proper soil fertility diagnosis, must be revised to optimize the benefits expected from inoculation including BNF. Starter N in the form of Vermicompost in low fertility soils at the rates used in this study did not suppress soybean nodulation, and it improved the productivity traits of the crop. However, further investigation is required to determine the threshold value of soil N content above which there will be no need to recommend starter N when rhizobia inoculants are applied to legume crops. This was beyond the scope of this study as many factors will have to be considered including soil types, mineralogy, weather conditions, legume genotype, rhizobia strains, and crop management.

## Acknowledgement

We are grateful for the financial support by the International Institute of Tropical Agriculture (IITA) under the COMPRO-II project funded by the Bill & Melinda Gates Foundation (OPPGD1398). We also acknowledge the contributions of Harrison Mburu, Elias Mwangi, Martin Kimanthi, and Philip Malala for their technical assistance during laboratory and greenhouse work.

---

**Fig 1:** Sites where the soils used in the first greenhouse experiment were collected including Trial Site 17 (i.e. Soil B) that was also used in both the second greenhouse experiment and the field trial, and Trial Site 7 (i.e. Soil A) used also in the field trial

**Fig 2:** Soybean nodule fresh weight and shoot dry weight in the first greenhouse experiment with and without co-application of Legumefix and Sympal (L+S) across sixty soils.

**Fig 3:** Soybean nodule fresh weight across trials following: (a) N-amendment in the form of vermicompost or urea co-applied with Legumefix and Sympal (L+S) in the second greenhouse trial (Soil B from Trial Site 17); (b) Legumefix application at Trial Site 7 and Trial Site 17 (field conditions); (c) Sympal application in field conditions; and (d) vermicompost in field conditions. The error bars represent the standard error of the difference (SED).

**Fig 4:** Soybean nodule effectiveness across trials following: (a) N-amendment in the form of vermicompost or urea co-applied with Legumefix and Sympal (L+S) in the second greenhouse trial (Soil B from Trial Site 17); (b) Legumefix and Sympal applications in field conditions at Trial Site 7 and Trial Site 17; and (c) vermicompost application in field conditions irrespective of the sites. The error bars represent the standard error of the difference (SEO).

**Fig 5:** Shoot dry weight across trials following: (a) N-amendment in the form ofvermicompost or urea co-applied with Legumefix and Sympal (L+S) in the second greenhouse trial (Soil B from Trial Site 17); (b) various combinations of vermicompost, Legumefix, and Sympal in field conditions; and (c) locations. The error bars represent the standard error of the difference (SED).

